# Widespread repurposing of the human CD38 ADP-ribosyl cyclase fold for toxicity in antibacterial and anti-eukaryotic bacterial polymorphic toxins

**DOI:** 10.1101/2025.07.23.666440

**Authors:** Julius Martinkus, Laurent Terradot, Dukas Jurėnas, Eric Cascales

**Author notes:** To whom correspondence should be addressed: Dukas Jurėnas, Eric Cascales.

## Abstract

Bacterial polymorphic toxins are modular weapons that mediate inter-microbial competition and host interactions by delivering diverse cytotoxic domains through specialized secretion systems. Here, we identify and characterize a novel toxin domain in *Pantoea ananatis* that displays remarkable structural and functional conservation with the human enzyme CD38. This bacterial toxin, fused to the type VI secretion system (T6SS) PAAR domain, harbors a C-terminal ADP-ribosyl cyclase (ARC) domain that hydrolyzes NAD^+^ and NADP^+^ *in vitro* and *in vivo*, leading to growth inhibition in both bacterial and eukaryotic cells. The 1.6-Å resolution structure of ARC reveals that it adopts a globular fold nearly identical to the human CD38 ADP ribosyl cyclase, with key catalytic residues conserved. ARC toxicity is neutralized in *P. ananatis* by a dedicated immunity protein. Comparative genomics reveals that CD38-like ARC domains are widespread in bacteria, fused to diverse delivery modules including T6SS, T7SS, and CDI systems. Functional assays demonstrate that these domains act as NAD-depleting toxins, with cross-immunity observed between non-cognate toxin–immunity pairs. Taken together, our findings reveal that a eukaryotic-like NAD^+^ hydrolase fold has been adapted in bacteria to generate a novel class of metabolic toxins, expanding the functional scope of polymorphic effectors and illustrating how conserved host-like enzymes can be co-opted for microbial warfare.

## INTRODUCTION

Bacteria live in densely populated, competitive environments where access to nutrients and space is limited (Little et al., 2008; Hibbing et al., 2010; Ghoul and Mitri, 2016; Garcia-Bayona and Comstock, 2018; Peterson et al., 2020). To thrive, many species have evolved potent weapons to inhibit or eliminate their rivals (Ruhe et al., 2013; Russell et al., 2014; Jamet and Nassif, 2015; Chassaing and Cascales, 2018; Klein et al., 2020; Kern et al., 2021; Cuthbert et al., 2022; Boardman et al., 2023). Among the most versatile of these weapons are polymorphic toxins (PT), which are modular, multidomain proteins that are delivered into neighboring cells to sabotage essential processes and cellular functions (Zhang et al., 2012; Jamet and Nassif, 2015; Ruhe et al., 2020). These toxins are often secreted through sophisticated nanomachines, such as the type IV (T4SS), type V (T5SS), type VI (T6SS) or type VII (T7SS) secretion systems (Shen et al., 2012; Souza et al., 2015; Willett et al., 2015; Bayer-Santos et al., 2019; Klein et al., 2020; Hernandez et al., 2020; Jurenas and Journet, 2021; Boardman et al., 2023; Lin, 2024), and are typically accompanied by immunity proteins that protect the producing bacterium from self-intoxication (Benz and Meinhardt, 2014; Jana and Salomon, 2019). The diversity and adaptability of polymorphic toxins have made them central players in interbacterial warfare, microbial community structuring, and even host interactions (Ruhe et al., 2020).

A defining feature of polymorphic toxins is their modularity. These proteins share a conserved N-terminal “cargo” or delivery domain fused to highly variable C-terminal “effector” domains, which confer toxic activity (Zhang et al., 2012; Jamet and Nassif, 2015; Ruhe et al., 2020). Over evolutionary time, this modularity has enabled the exchange, shuffling, and specialization of toxin domains, resulting in a vast repertoire of enzymatic activities including nucleases, peptidases, lipases, or NAD-targeting enzymes (Zhang et al., 2012). Increasingly, it has become clear that bacteria do not solely evolve new toxic domains de novo; they also select, co-opt and repurpose host-like proteins for bacterial warfare (Ruhe et al., 2020; Aravind et al., 2024; Anshul and Kumari, 2025).

One of the most intriguing examples of repurposing is the recruitment of eukaryotic-like domains into bacterial toxins, a process that blurs the lines between pathogenicity and mimicry (Chen and Xia, 2021; Martyn et al., 2022; Anshul and Kumari, 2025). This phenomenon reflects a broader evolutionary strategy in which bacteria hijack host-derived folds and functions, modifying them for use in competition, colonization, or immune evasion (Chen and Xia, 2021; Martyn et al., 2022). Notably, several bacterial toxins appear to mimic or structurally resemble eukaryotic proteins involved in key cellular processes, such as actin polymerization, cell signaling, and nucleotide metabolism (Stebbins and Galan, 2001; Mak and Thurston, 2021, Chen and Xia, 2021; Anshul and Kumari, 2025).

In this study, we describe the discovery of a novel toxin domain in *Pantoea ananatis* that exemplifies this strategy of molecular hijacking. The toxin, encoded within a T6SS *vgrG*/*hcp*/*paar* island, harbors an N-terminal Proline-Alanine-Alanine-Arginine (PAAR) cargo domain, putative transmembrane helices and a C-terminal domain with strong structural and functional similarity to the human enzyme CD38 (Cluster of Differentiation 38), a multifunctional ADP-ribosyl cyclase (ARC) involved in immune cell activation and regulation of calcium intracellular levels and signaling (De Flora et al., 1997; Schuber and Lund, 2004; Lee, 2012; Morandi et al., 2019; Manik et al., 2022). We show that this bacterial toxin, designated ARC^tox^, shares the catalytic residues, enzymatic activity, and overall fold of CD38, yet functions as a cytotoxic effector capable of depleting NAD^+^ and NADP^+^ in both bacteria and eukaryotic target cells, suggesting a dual role in microbial competition and host interaction. By tracing the distribution of ARC-like domains across bacterial genomes, we reveal that CD38-like folds have been extensively co-opted by bacteria and fused to diverse secretion and delivery systems. This widespread appropriation of a eukaryotic enzyme scaffold underscores the remarkable evolutionary plasticity of PT, and positions ARC^tox^ as a model for understanding how metabolic enzymes can be transformed into toxic effectors.

Together, our findings highlight that domestication of host-like metabolic enzymes into modular toxins is an efficient strategy in bacterial warfare. This not only expands the known functional repertoire of PT but also deepens our understanding of how bacteria evolve through innovation-by-mimicry, repurposing host or bacterial proteins for competitive advantage.

## RESULTS

### The C-terminal domain of the *Pantoea ananatis* PAAR PANA_2924 protein is toxic when expressed in the *E. coli* or yeast cytosol

The plant pathogen *Pantoea ananatis* strains LMG 2665^T^ and DZ-12 use a type VI secretion system (T6SS) to promote virulence in onion plants and to outcompete various bacterial species, including *Escherichia coli* (Shyntum et al., 2015; Zhao et al., 2024). Other *Pantoea* species encode up to three distinct T6SS gene clusters, each containing all essential structural components, as well as a variety of *vgrG*, *hcp*, and *PAAR* islands (de Maayer et al., 2011; Shyntum et al., 2014; Weller-Stuart et al., 2017). In addition to T6SS needle components, these genomic islands often encode effector proteins, including the recently characterized TseG nuclease (Zhao et al., 2024).

In the *P. ananatis* reference strain LMG 20103, one such *PAAR* island comprises three genes encoding a putative EagR-family chaperone (PANA_2925), which usually binds to and protects effector transmembrane domains (Alcoforado Diniz *et al*., 2015; Quentin *et al*., 2018; Ahmad *et al*., 2020; Jurenas *et al*., 2021), a polymorphic PAAR protein with an extended C-terminal region of unknown function (PANA_2924), and a small protein (PANA_2923) (Figure 1A). This gene organization resembles canonical chaperone-dependent polymorphic toxin operons, where a central toxin gene fused to a cargo is flanked by a chaperone and an immunity protein. Structural predictions and sequence alignments of PANA_2924 revealed three distinct domains: a canonical PAAR domain (residues 1–21 and 72–168), which could associate with the T6SS needle spike, a five-helix bundle (residues 22–71 and 169–249), and a globular C-terminal extension (residues 255–387) (Figures 1B and 1C). It is noteworthy that the PAAR domain is interrupted by two helices from the central bundle, suggesting a structurally integrated architecture (Figure 1B).

**Figure 1.**
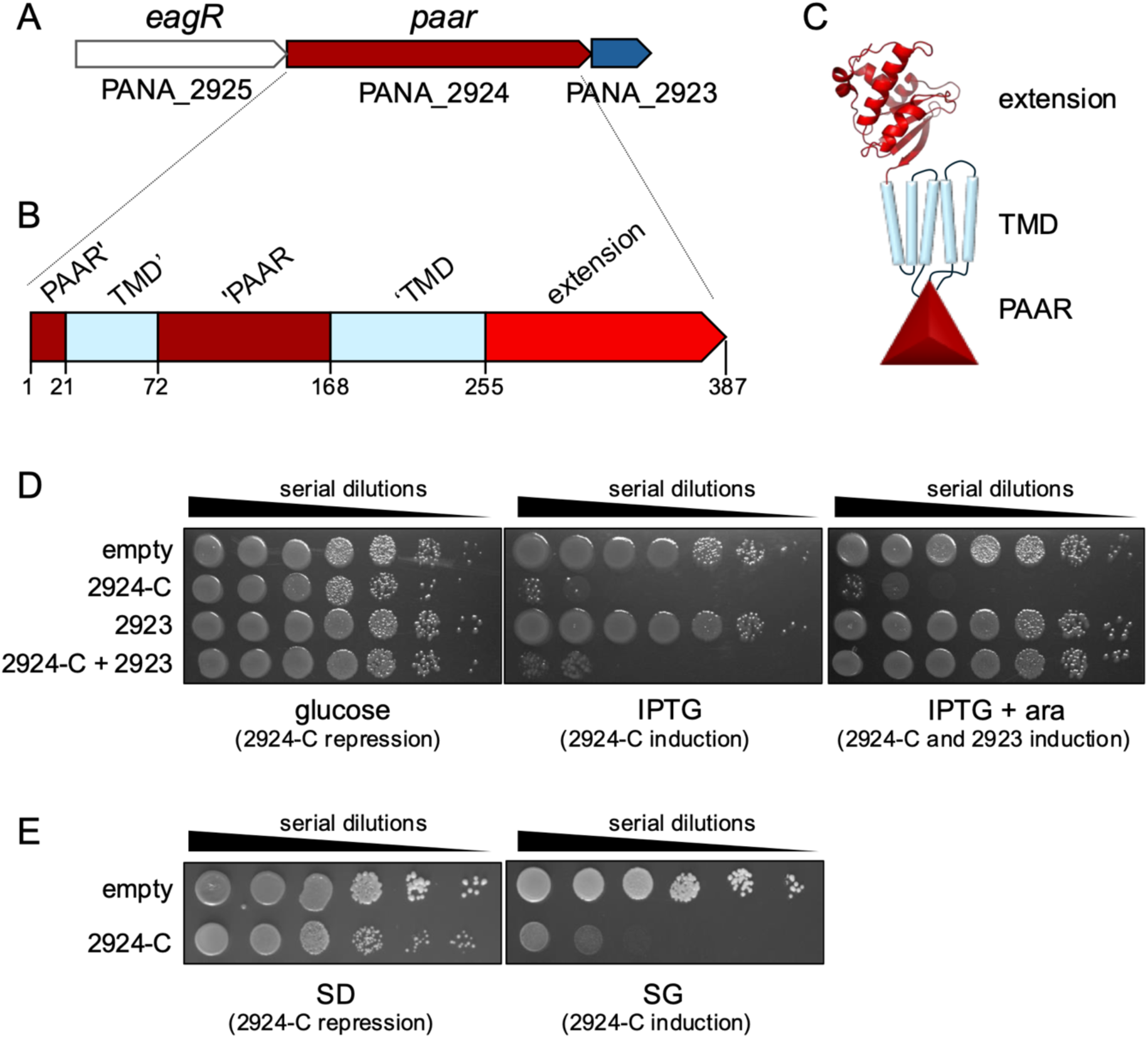
*Pantoea ananatis* PANA_2024 C-terminal domain is toxic in *E. coli* and yeast cells. **(A)** Schematic representation of the *P. ananatis PANA_2925-PANA_2023* PAAR island. Genes encoding the EagR chaperone (PANA_2925), the specialized PAAR protein (PANA_2924), and a protein of unknown function (PANA_2923) are depicted in white, red and blue, respectively. **(B and C)** Schematic representation of the specialized PAAR protein, highlighting the different domains and their boundaries: PAAR domain (dark red), trans-membrane domain (TMD, light blue) and C-terminal extension (red). **(D)** Toxicity assay in the heterologous host *E. coli.* Overnight cultures of *E. coli* cells expressing the PANA_2924 C-terminal extension (2924-C) from the low-copy vector pNDM220, PANA_2923 (2923) from the pBAD33 vector, or both (2924-C+2923) were serially diluted (10^-1^ to 10^-6^) and spotted on LB agar plates, supplemented with 1% of glucose (left panel, 2924-C repression), with 0.05 mM of IPTG (middle panel, 2924-C induction), or with 0.05 mM of IPTG and 1 % of L-arabinose (right panel, 2924-C and 2923 induction). **(E)** Toxicity assay in yeast. Overnight cultures of W303 cells expressing 2924-C from the pRS416_Gal1 vector were serially diluted and spotted on SD (left panel, 2924-C repression) or SG (right panel, 2924-C induction) medium.

To test whether the C-terminal extension corresponds to a toxin module, its sequence was cloned under the control of the P*_lac_* promoter into the low-copy pNDM220 vector, and its toxicity was tested in *E. coli*. While the domain was not toxic when its expression was repressed by the presence of glucose, induction with IPTG led to strong growth inhibition, indicating antibacterial activity in the *E. coli* cytosol (Figure 1D). The *P. ananatis* T6SS being involved in virulence towards onions (Shynthum *et al*., 2015; Zhao *et al*., 2024), we tested whether the C-terminal extension of the specialized PAAR protein was also toxic into eukaryotic cells. The same domain was expressed in *Saccharomyces cerevisiae* from the single-copy pRS416 plasmid. Yeast growth was also inhibited (Figure 1E), demonstrating that the C-terminal extension of the PANA_2924 specialized PAAR protein has dual antibacterial and anti-eukaryotic activity.

### PANA_2923 encodes an immunity protein that neutralizes PANA_2924 toxicity through protein-protein interaction

To determine whether the downstream gene, PANA_2923, encodes a cognate immunity protein conferring protection against PANA_2924, it was cloned under the *ParaBAD* promoter in the pBAD33 vector and co-expressed with the toxic PANA_2924 C-terminal domain. Co-expression fully rescued *E. coli* growth (Figure 1D), demonstrating that PANA_2923 neutralizes the toxin’s activity.

Co-purification assays confirmed a direct interaction between untagged PANA_2923 and 6×His-TEV-tagged PANA_2924 C-terminal domain (Figure 2A). Size exclusion chromatography further suggests a 1:1 stoichiometry of the complex (Figure 2B), consistent with immunity proteins binding to and blocking effector active sites.

**Figure 2.**
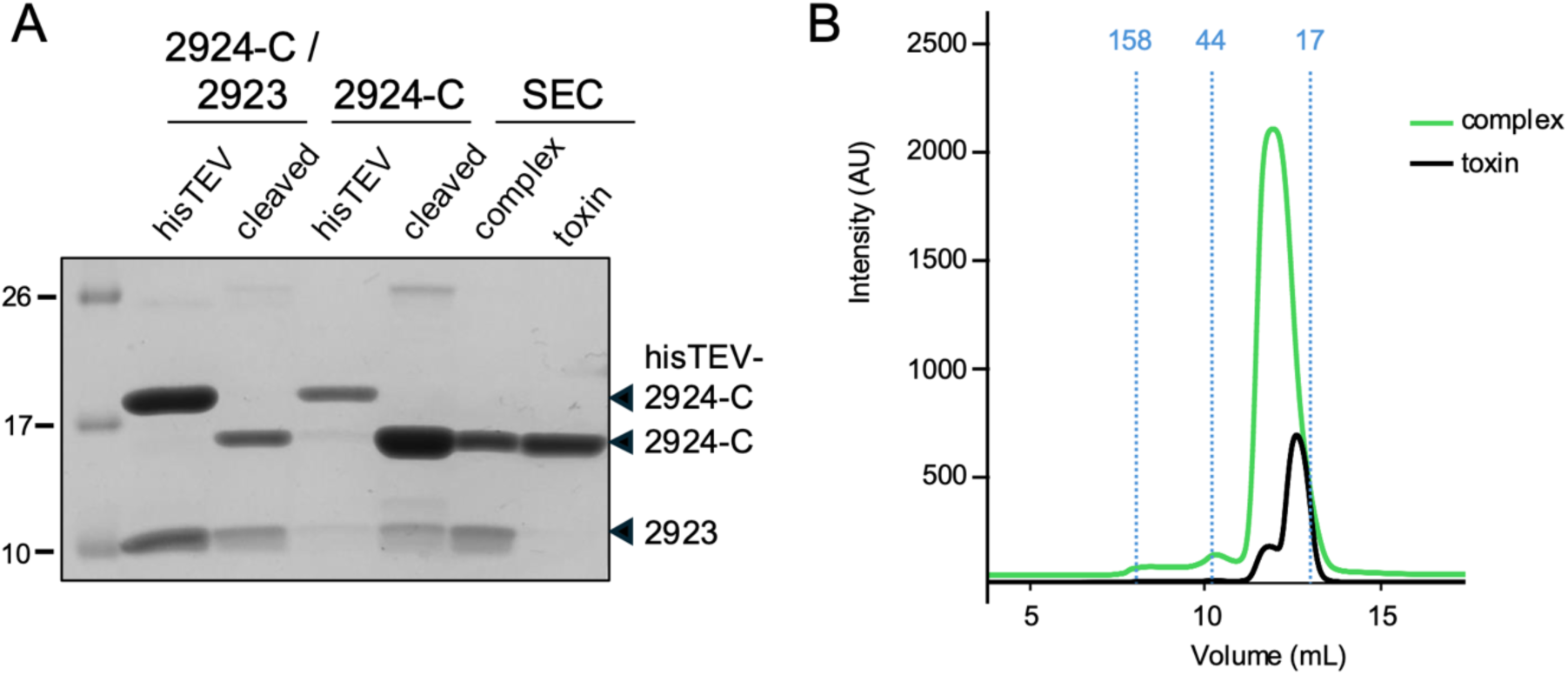
PANA_2924 C-terminal extension forms a complex with PANA_2923. **(A)** SDS-PAGE (12.5% acrylamide) and Coomassie blue staining of purified fractions: complex after affinity purification (lane 1), complex after TEV digestion (lane 2), refolded toxin before TEV digestion (lane 3), toxin after TEV digestion (lane 4), complex (lane 5), and toxin (lane 6) after TEV digestion and size-exclusion chromatography (SEC). The positions of the proteins are indicated on the right. Molecular weight markers (in kDa) are indicated on the left. **(B)** Gel filtration analysis. The PANA_2924-C / PANA_2923 complex (green line) and the PANA_2924-C toxin (black line) obtained after affinity purification and TEV cleavage were subjected to size-exclusion chromotography on a Superdex 75 10/30 column. Molecular mass standards (blue dotted lines) are indicated at the top (in kDa).

### PANA_2924 C-terminal domain structure reveals an ADP-ribosyl cyclase fold

The purified complex between the PANA_2924 C-terminal domain and its PANA_2923 immunity was subjected to crystallization trials, without success. Crystallization of the PANA_2924 extension was however achieved after dissociating the immunity protein from the complex by adding urea in the affinity chromatography washing buffer, followed by refolding and SEC purification of the isolated toxin. A diffraction dataset was collected and the structure was solved by molecular replacement using the AlphaFold3 model and refined to a resolution of 1.6 Å (Figure 3A, Supplemental Table S1). PANA_2924 crystallized in the space group P2_1_2_1_2_1_, with two molecules per asymmetric unit, which adopt similar conformations: while the first chain could be modeled from residues 252 to 385, missing a flexible loop from 311 to 317, the second chain could be modelled from residues 250 to 385, missing a different flexible loop (amino-acids 345 to 349) (Supplementary Figure S1). The structure reveals a globular fold with two lobes, consisting of a four-stranded β-sheet covered by a helical cap composed of five α-helices (Figure 3A). A search for structural homologues using DALI (Holm, 2022) showed that the PANA_2924 C-terminal extension displays a fold similar to the human protein CD38 (Figure 3B). CD38 is a multifunctional protein: CD38 is a receptor at the surface of immune cells, responsible for the activation of T cells for the production of cytokines. CD38 has also ADP ribosyl cyclase (ARC) activity: it hydrolyzes NAD^+^ to generate cyclic ADP ribose (cADPr), a messenger mediating various physiological functions in eukaryotic (notably involved in calcium signaling) and bacterial cells (Schuber and Lund, 2004; Lee, 2012; Ferrero et al., 2014; Takasawa, 2022; Lee et al., 2022). Importantly, key catalytic residues of CD38 – two glutamates and two tryptophans located in the cavity between the β-sheet and the α-helical cap (Munshi et al., 1999; Yamamoto-Katayama et al., 2002; Schuber and Lund, 2004; Kuhn et al., 2014) – are conserved in the PANA_2924 ARC toxin (hereafter named ARC^tox^) (Figure 3C). These residues (W33, E59, W83, E116) form a putative active site. In addition, the AlphaFold3 model of the ARC^tox^ with its immunity protein PANA_2923 (ARC^imm^) suggests that ARC^imm^ interacts extensively with the ARC^tox^ catalytic cleft (interface area of 1130 Å), with a loop projecting into the active site (Figures 3D and 3E). In agreement with this information, substitution of any of the ARC^tox^ putative catalytic residues, W33, E59, W83 and E116, by alanines abolished ARC^tox^ activity in *E. coli* (Figure 3F).

**Figure 3.**
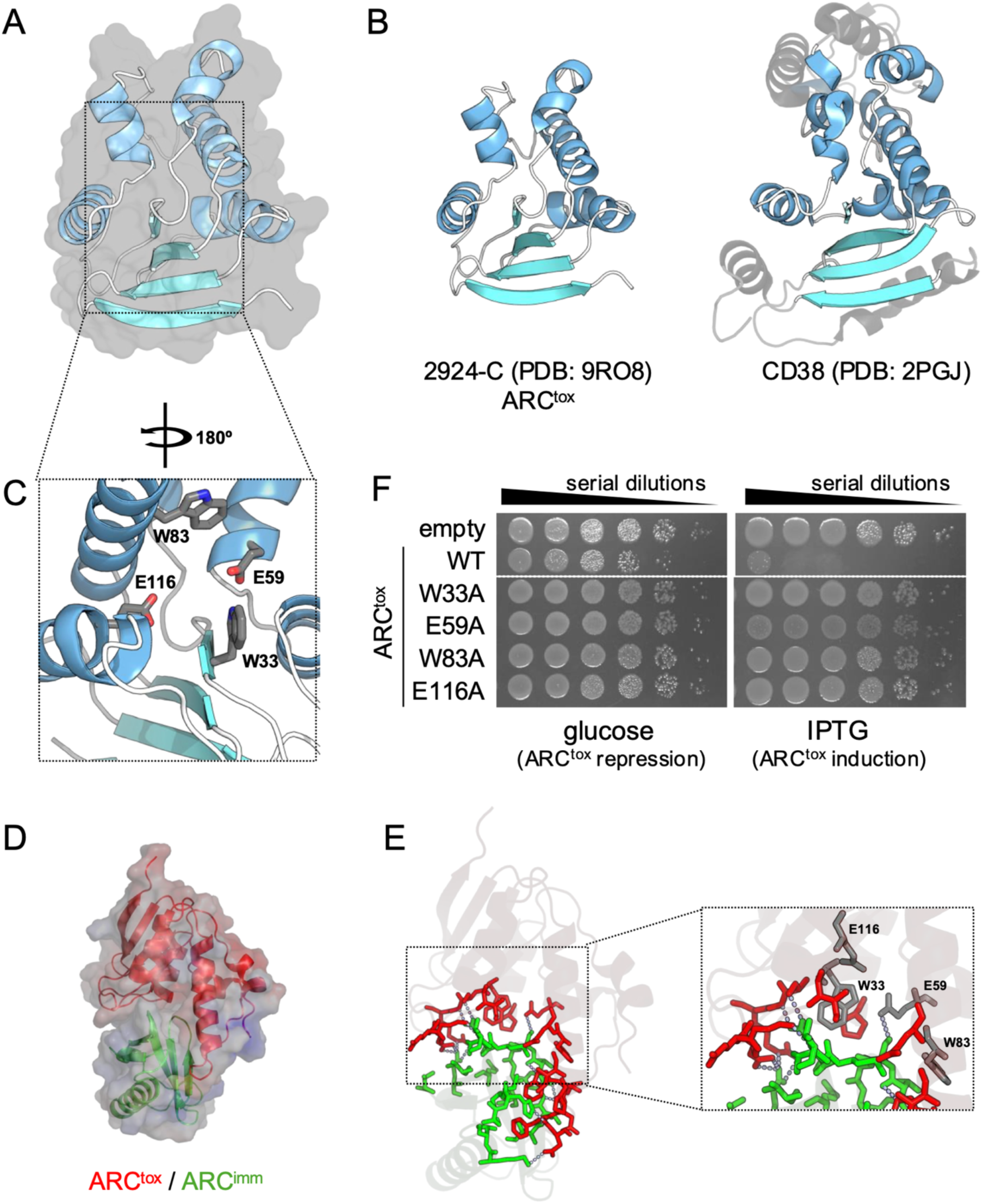
PANA_2924 C-terminal extension displays an ADP ribosyl cyclase (ARC) fold comparable to human CD38, with a typical catalytic cleft blocked by the PANA_2923 immunity protein. **(A)** Ribbon representation of the 1.6-Å resolution crystal structure of PANA_2924 C-terminal extension. α-helices are depicted in light blue whereas β-strands are shown in cyan. The surface structure is shown in grey transparent. **(B)** Structural comparison between *P. ananatis* PANA_2924 C-terminal extension (this work) and human CD38 protein (PDB: 2PGJ). Conserved secondary structures are colored as in panel **A**. **(C)** Magnification of the active site, showing putative key residues as grey sticks. **(D)** Ribbon representation of the AlphaFold3 prediction model of the ARC^tox^/ARC^imm^ complex (ipTM score = 0.92). The toxin domain is shown in red whereas the immunity is shown in green within the complex surface representation colored based on residue electrostatic properties (from blue (positive) to red (negative)). **(E)** Contact site between ARC^tox^ and ARC^imm^ highlighting the ARC^imm^ loop penetrating into the ARC^tox^ catalytic cleft. Potential electrostatic and van der Waals interactions are depicted by dotted grey lines. **(F)** Toxicity assay in the heterologous host *E. coli.* Overnight cultures of *E. coli* cells expressing the wild-type (WT) or mutated ARC^tox^ from the low-copy vector pNDM220 were serially diluted (10^-1^ to 10^-6^) and spotted on LB agar plates, supplemented with 1% of glucose (left panel, ARC^tox^ repression), or 0.05 mM of IPTG (right panel, ARC^tox^ induction).

### ARC^tox^ hydrolyzes NAD^+^ and NADP^+^ *in vitro* and *in vivo* but does not produce cADPr

We next assessed the enzymatic activity of the purified, refolded, ARC^tox^ domain. *In vitro* assays showed efficient degradation of both NAD^+^ and NADP^+^ (Figure 4A). NAD^+^ degradation activity was confirmed *in vivo*, by measuring total NAD^+^ levels in cells after 1 hour of induction of the ARC^tox^ domain: Figure 4B shows that expression of ARC^tox^ led to a drastic decrease of cellular NAD^+^ levels, confirming enzymatic activity in the cytosol. Co-expression of ARC^imm^ or expression of the ARC^tox^ catalytically inactive mutants restored NAD^+^ levels to near baseline, except for the E59A mutant, which retained a level of degradation comparable to that of the wild-type toxin.

**Figure 4.**
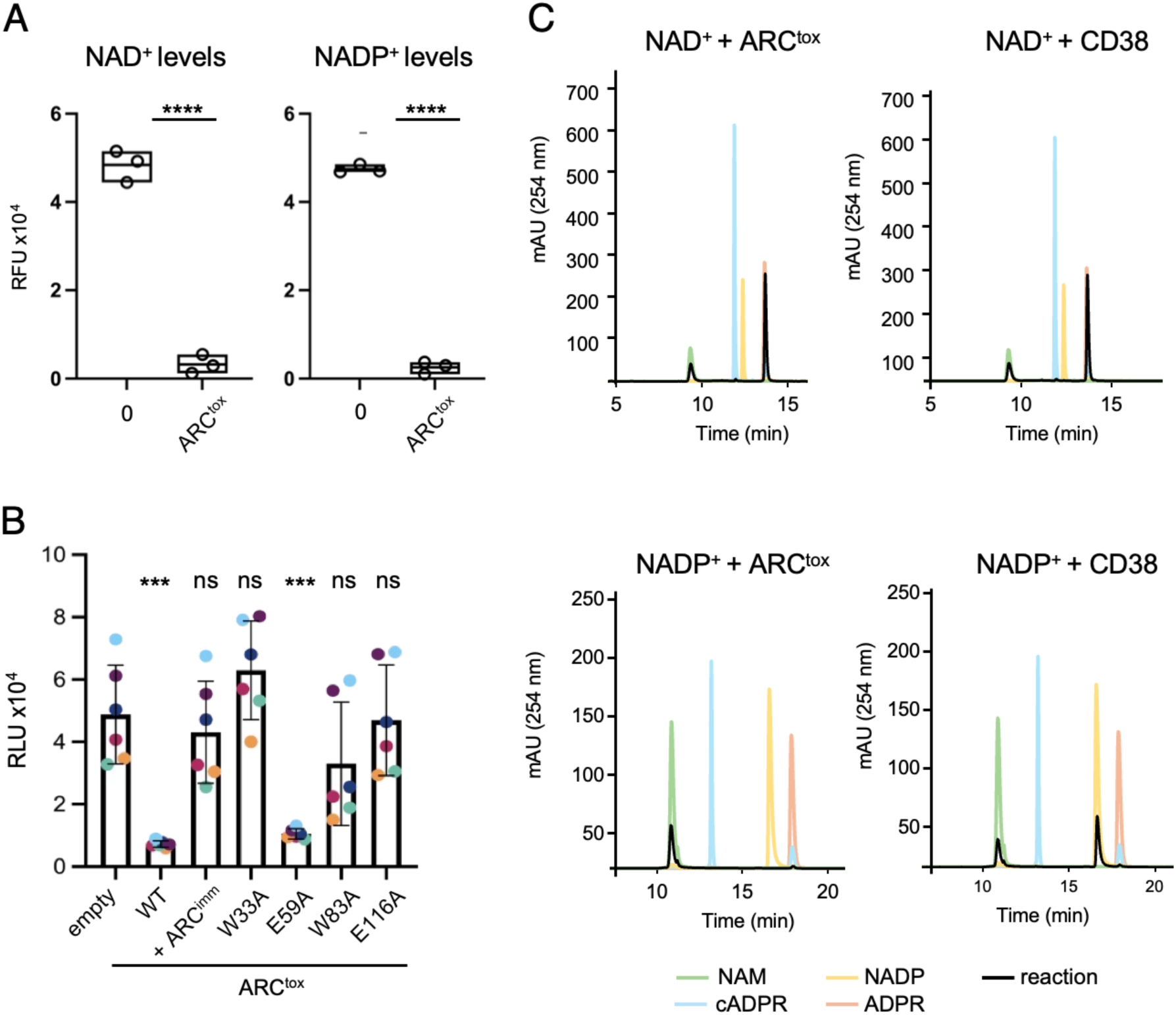
*P. ananatis* ARC^tox^ depletes NAD^+^ and NADP^+^ pools *in vivo* and *in vitro*. **(A)** Fluorescence-based assay showing NAD^+^ and NADP^+^ levels in untreated (0) and ARC^tox^-treated samples. NAD^+^ (left panel) or NADP^+^ (right panel) (0.6 mM) were incubated with the buffer (0) or with refolded ARC^tox^ (0.1 μM) *in vitro* for 30 min. RFU – relative fluorescence units. Boxes indicate standard deviation of the mean of three replicates. **** denotes a statistically significant difference between the two condiutions, based on an unpaired t-test (P<0.0001). **(B)** Luminescence assay measuring NAD^+^/NADH levels in *E. coli* DH5α cells producing the wild-type (WT) or the indicated mutated ARC^tox^ domains from the pNDM220 vector and ARC^imm^ from the pBAD33 vector. RLU – relative luminescence units. Bars show means ± SD from six independent experiments (same-color points represent replicates from the same experiment). *** denotes statistically significant difference compared to the control group (one-way ANOVA, post-hoc Dunnett’s test, P<0.001). ns = not significant. **(C)** HPLC chromatograms of the products of the reactions of NAD^+^ (upper panels) and NADP^+^ (lower panels) in the presence of ARC^tox^ (left panels) or CD38 (right panels) subjected to reverse-phase (RP) Atlantis Premier BEH C18 AX column. The products are shown with the black lines. Chomatograms of selected standard analytes are shown in color (green, nicotinamide; orange, NADP; pink, ADP ribose; blue, cyclic ADP ribose).

To get further information on the reaction catalyzed by ARC^tox^, we performed high-pressure liquid chromatography (HPLC) to identify the reaction products. Upon incubation with ARC^tox^, NAD^+^ and NADP^+^ were converted into nicotinamide and ADP ribose (Figure 4C panels 1 and 3). No cADPr was however detected, likely due to its immediate hydrolysis to ADP ribose (Sauve et al., 1998). Consistent results were obtained with the well-characterized human CD38 ADP ribosyl cyclase (Figure 4C, panels 2 and 4), further supporting functional conservation.

### ARC domains are widespread and associated with polymorphic toxin systems

To explore the evolutionary context of ARC^tox^, we searched for homologous domains across bacterial genomes. ARC-like domains were found throughout the bacterial tree of life (Figures 5A and 5B), exclusively associated with secretion-targeting domains such as PAAR, RHS, FIX (T6SS), LXG and EsxA (T7SS), TPS (two-partner T5SS), and other extracellular or cell-wall anchoring elements like pyocins, Flp pilins, and fibronectin type III (Fn3) domains (Schneider et al., 2013; Whitny et al., 2017; Jana et al., 2019; Wood et al., 2019; Jurenas et al., 2021)(Figure 5A). Structural modeling showed that all these ARC homologs adopt the typical CD38-like ADP ribosyl cyclase fold (Supplementary Figure S2).

**Figure 5.**
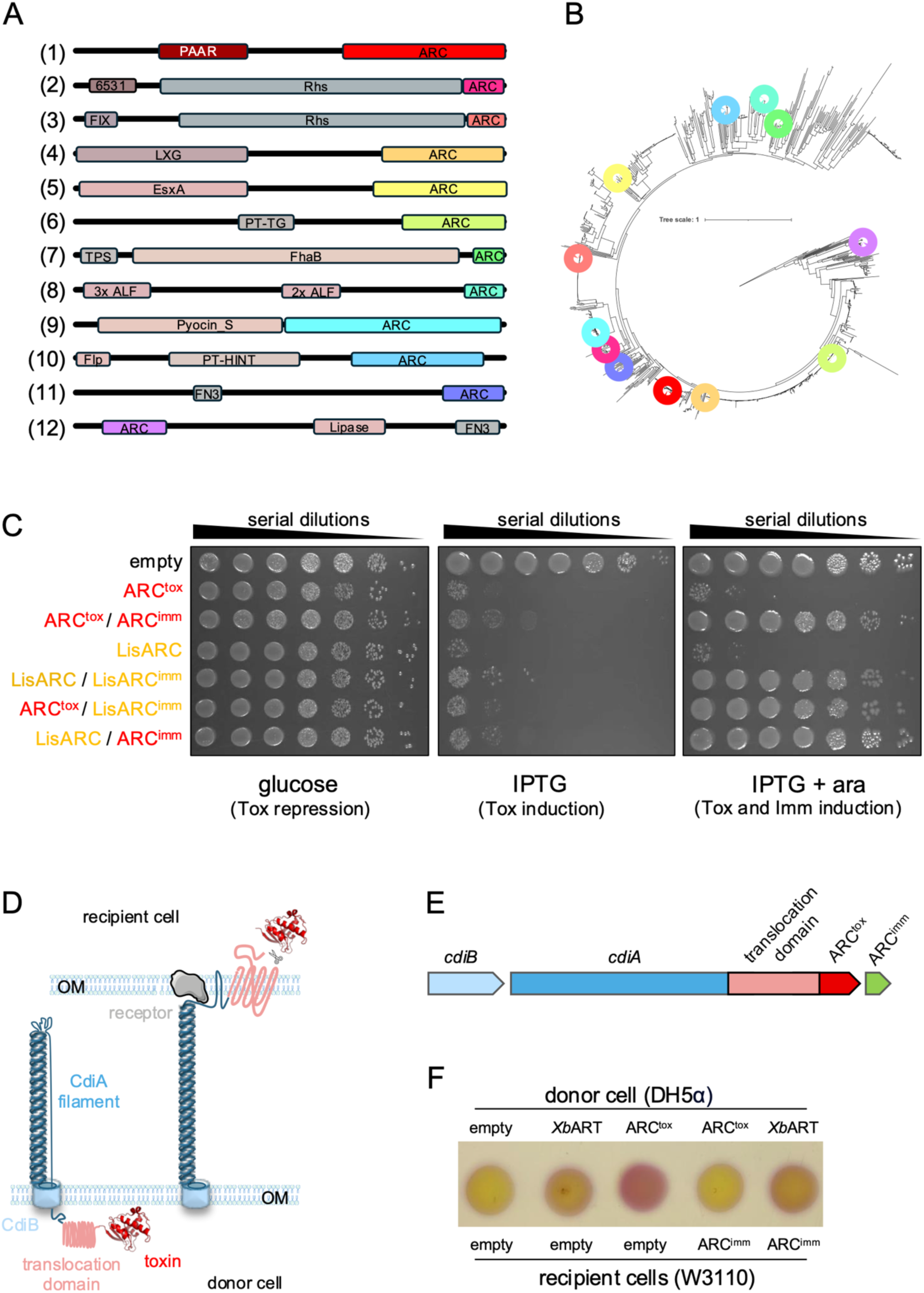
Active ARC toxin domains are widespread across bacteria and can be used for engineering. **(A)** Domain architectures of selected proteins containing ARC domains homologs (1, *Pantoea ananatis* WP_013026797; 2, *Agrobacterium rubi* NTF28035; 3, *Burkholderia gladioli* WP_186165364; 4, *Listeria monocytogenes* WP_120135998; 5, *Gordonia jinhuaensis* WP_188589035; *Lactiplantibacillus plantarum* WP_076633504; 7, *Stenotrophomonas maltophilia* OCK46403; 8, *Nonomuraea fuscirosea* WP_364663299; 9, *Marinobacter shengliensis* WP_138437386; 10, *Pendulispora albinea* WP_394825747; 11, *Allomuricauda sp.* RPG31737; 12, *Streptomyces mirabilis* WP_388533272). ARC domains are indicated in bright colors **(B)** Cladogram of 1075 homologs of *P. ananatis* ART^tox^. The positions of the selected ARC domains shown in panel A are indicated by the same color code. **(C)** Toxicity assay in the heterologous host *E. coli.* Overnight cultures of *E. coli* cells expressing the indicated ARC domain (*P. ananatis* ARC^tox^ or *L monocytogenes* ARC, LisARC) from the low-copy vector pNDM220, and the indicated immunity proteins (*P. ananatis* ARCimm and *L. monocytogenes* LisARC^imm^) from the pBAD33 vector were serially diluted (10^-0^ to 10^-6^) and spotted on LB agar plates, supplemented with 1% of glucose (left panel, toxin repression), 0.02 mM of IPTG (middle pannel, toxin induction), and 0.02 mM of IPTG and 1% of arabinose (right panel, toxin and immunity induction). **(D)** Schematic model of Contact-Dependent Inhibition (CDI) system mechanism of action. The outer membrane component CdiB translocates CdiA (blue) to the donor cell surface. The receptor-binding domain of CdiA recognizes and binds to a specific receptor (grey) on the surface of the recipient cell. Then the translocon domain (pink) delivers the toxin (red) into the recipient cell. **(E)** Schematic representation of the CDI chimera engineered for ARC^tox^ delivery by the *E. coli* CdiAB CDI system. The *cdiB* and *cdiA* genes are shown in light blue and blue, respectively. The grafted translocation domain and *P. ananatis* ARC^tox^ domain are shown in light red and red, respectively. ARC^imm^ is shown in green. **(F)** Interbacterial competition assay. Donor (*E. coli* DH5α) and recipient (*E. coli* W3110) cells producing the indicated chimera CDIs (*P. ananatis* ARC^tox^ and *Xenorhabdus bovieni* ART) or ARC^imm^ were mixed at a 1:2 (v:v) ratio, spotted onto LB-agar, and incubated for 4 h at 37°C. After competition recipient cell lysis was revealed using CPRG (yellow), which turns purple upon hydrolysis by β-galactosidase from lysed *lacZ^+^* recipient cells.

Based on amino-acid sequence identity (>70%), the closest homologues of *P. ananatis* ARC^tox^ are encoded within *Listeria* spp genomes and systematically associated with T7SS-dependent LXG domains. Toxicity assays of a representative T7SS ARC domain from *Listeria monocytogenes* (LisARC) confirmed that it is toxic when expressed in *E. coli* and was neutralized by its cognate immunity protein (Figure 5C). In agreement with the high identity between ARC^tox^ and LisARC, cross-protection experiments demonstrated that *Pantoea* ARC^tox^ and LisARC are both neutralized by non-cognate immunity proteins (Figure 5C), suggesting recent recruitment of ARC domains to different secretion systems.

### ARC^tox^ can be secreted via a heterologous delivery system

To test whether ARC^tox^ could be redirected through another delivery system, we engineered a chimera using the *E. coli* CdiAB contact-dependent inhibition (CDI) system, as done recently with the *Xenorhabdus bovieni* TreX ADP-ribosyl transferase (*Xb*ART, Dumont et al., 2024). For this, ARC^tox^ was fused to the C-terminal CdiA stick domain, after the conserved VENN motif that delimitates CdiA transport and toxic domains (Figures 5D and 5E). Exposition of this chimeric effector to the *E. coli* DH5α cell surface was toxic to *E. coli* W3110 competitor cells but not to cells expressing the ARC immunity protein, demonstrating effective delivery and functional toxin activity (Figure 5F). In contrast, ARC^imm^ did not protect against an unrelated CDI toxin (*Xb*ART), indicating specific poisoning of recipient cells with ARC^tox^ (Figure 5F).

## DISCUSSION

In this manuscript, we report the characterization of a novel and conserved class of versatile polymorphic toxin domains, ARC^tox^, sharing structural and functional characteristics of the human CD38 ADP ribosyl cyclase. We demonstrate that the C-terminal extension of a specialized PAAR protein from *Pantoea ananatis*, PANA_2924, acts as a potent cytotoxic effector, exhibiting both antibacterial and anti-eukaryotic activity. This toxin is neutralized by a tightly binding immunity protein (PANA_2923), forming a canonical toxin–immunity module reminiscent of those employed in interbacterial conflict. Very recently, the antibacterial Tac1 toxin associated with the *Streptoccocus parasanguinis* T7SS was functionally and structurally characterized, demonstrating that it rapidly hydrolyzes NAD^+^ and NADP^+^ (12,000 molecules/min) into ADP-ribose and ADP-ribose 2’-phosphate, respectively (Colautti et al., 2025).

Remarkably, structural and biochemical data indicate that ARC^tox^ and Tac1 are bacterial analog of human CD38, a multifunctional enzyme involved in immune signaling and NAD metabolism (Lee et al., 2022; Takasawa, 2022; Colautti et al., 2025), revealing an unexpected link between polymorphic bacterial effectors and eukaryotic cell biology. The discovery of this relationship between ARC^tox^ and Tac1 and CD38 is both structurally and functionally significant. The ARC toxins adopt a globular fold closely resembling CD38, characterized by a β-sheet core capped by α-helices. DALI analysis and high-resolution crystal structures reveal that these enzymes share not only overall architecture but also precise placement of key catalytic residues – two glutamates and two tryptophans – which define the active site of the CD38 family (Munshi et al., 1999; Schuber and Lund, 2004; Kuhn et al., 2014). Mutational inactivation of these residues in ARC^tox^ completely abrogates its toxic activity, confirming their critical role in catalysis. Importantly, this enzymatic function is not merely structural mimicry: ARC^tox^ directly hydrolyzes NAD^+^ and NADP^+^ into ADP-ribose and nicotinamide, recapitulating the enzymatic outcome of CD38 activity in eukaryotic cells (Schuber and Lund, 2004). The absence of cyclic ADP-ribose (cADPr), likely due to its rapid hydrolysis, aligns with the catalytic profile of some CD38-like enzymes that favor hydrolysis over cyclization (Sauve et al., 1998). These data also suggest that ARC^tox^ antibacterial toxicity is not due to the accumulation of cADPr, which could impact undefined signaling pathway in bacteria, but rather poisons target cells by depleting the pool of NAD^+^/NADP^+^. Indeed, the recent work of Colautti et al. demonstrated that the *S. parasanguinis* T7SS Tac1 ARC toxin hydrolyzes NAD^+^ with a rate of 12,000 molecules/min (Colautti et al., 2025), which is comparable with previous NAD glycohydrolase toxins (Whitney et al., 2015; Tang et al., 2018). The ability of a bacterial toxin to deplete NAD^+^, a central metabolic cofactor, highlights a sophisticated mechanism of intercellular antagonism that targets core bioenergetic processes. While many known bacterial toxins act by degrading nucleic acids, lipids or proteins (Zhang et al., 2012; Russell et al., 2014; Durand et al., 2014; Jurenas and Journet, 2021), ARC^tox^ represents a distinct class of metabolic toxins, sabotaging host or competitor metabolism through NAD^+^ depletion. Other toxins belonging to the ART-fold superfamily have been shown to utilize the NAD^+^ or NADP^+^ for ADP-ribosylation of protein or of RNA targets, or for NAD^+^ hydrolysis (Suarez et al., 2010; Whitney et al., 2015; Palazzo et al., 2017; Ting et al., 2018; Tang et al., 2018; Jurenas et al., 2021b; Roussin and Salcedo, 2021; Jurenas et al., 2022; Bullen et al., 2022). This ART-fold superfamily comprises a conserved core fold corresponding to a central split β-sheet with a conserved order of strands (Aravind et al., 2015). The ARC fold is highly distinct, but our genomic analysis showed that it is also highly widespread, but previously only reported in eukaryotic CD38. In any case, the NAD^+^ depletion strategy likely confers a competitive advantage in polymicrobial environments, especially in the plant rhizosphere or during host colonization, where nutrient access and niche competition are intense (Hibbing et al., 2010; Ghoul and Mitri, 2016; Venturi and Keel, 2016). The dual activity of ARC^tox^ against both bacterial and eukaryotic cells further suggests that this effector may function during both interbacterial competition and plant–pathogen interactions, consistent with the known virulence roles of *P. ananatis* T6SS in onion infection (Shyntum et al., 2015; Zhao et al., 2024).

The conservation between ARC^tox^ and CD38 extends beyond structure and catalysis; it raises fascinating evolutionary questions. The phylogenetic distribution of ARC domains across diverse bacterial lineages, and their consistent association with modular toxin architectures (PAAR, RHS, LXG, TPS, Fn3, etc.), suggests that these domains have been evolutionarily co-opted for secretion via multiple delivery systems, including T6SS, T5SS, T7SS, and CDI, or for cell surface exposition. This modularity, which is a hallmark of polymorphic toxins, enables effector domains to be flexibly redeployed across different antagonistic contexts. Yet the fact that the same enzymatic fold underlies both a eukaryotic signaling enzyme and a prokaryotic toxin raises the possibility of an ancient common ancestry or repeated events of horizontal gene transfer followed by domain repurposing. The metabolic versatility of the CD38/ARC fold, capable of catalyzing both cyclization and hydrolysis reactions on NAD^+^ substrates, may have facilitated its functional diversification across kingdoms.

Our findings also underscore the high adaptability of bacterial effector systems. The incorporation of ARC^tox^ into a heterologous CDI system, where it successfully mediates contact-dependent killing of target cells, highlights the modularity and portability of these effectors. This property, combined with the cross-reactivity of immunity proteins (i.e., cross-protection between *Pantoea* and *Listeria* ARC^tox^ modules), suggests that ARC^tox^ domains can be rationally engineered for synthetic biology applications, such as programmable antibacterial agents, targeted delivery systems, or novel biocontrol strategies in agriculture. Moreover, the cross-protection between non-cognate immunity proteins may reflect conserved structural features of the catalytic cleft, hinting at convergent evolution or selective pressure to maintain compatibility within toxin-immunity networks.

The biological implications of ARC^tox^ extend to host-pathogen interactions as well. Given its demonstrated anti-eukaryotic activity in yeast, one may hypothesize that ARC^tox^-like proteins are deployed during plant infection, where they could modulate host immunity, disrupt cellular metabolism, or interfere with NAD^+^-dependent signaling pathways. Such roles would mirror the function of CD38 in immune cells, albeit in a pathogenic rather than physiological context. Further investigations into the role of ARC^tox^ in plant–microbe interactions, including its potential delivery into plant cells or modulation of host NAD^+^ pools, could reveal novel mechanisms of bacterial virulence.

Our broader genomic survey of ARC domain distribution indicates that this ARC NAD hydrolase toxin architecture is widespread across the bacterial tree of life. Its association with diverse secretion and anchoring domains supports the notion that ARC^tox^ is a core component of the bacterial competitive arsenal. The presence of ARC domains in systems as distinct as pyocins and T7SS suggests that ARC^tox^ may contribute to antagonism in a variety of ecological contexts, from soil and plant microbiomes to animal hosts and clinical infections.

## EXPERIMENTAL PROCEDURES

### Strains, media and growth conditions

*Escherichia coli* strain DH5α was used for cloning, toxicity tests, and NAD/NADH-Glo^TM^ Assay (Promega). *E. coli* BL21 (DE3) and W3110 strains were used for protein production and bacterial competition, respectively. *E. coli* cells were grown at 37°C in Lysogeny Broth (LB) with agitation or on LB agar (1.5%) plates. When needed, media were supplemented with ampicillin (50-100 μg.mL^-1^), chloramphenicol (30 μg.mL^-1^), kanamycin (50 μ g.mL^-1^), or streptomycin (100 μ g.mL^-1^). Gene expression was induced by the addition of 1% of L-arabinose, 50-100 μM of IPTG or repressed with 1% of glucose. *Saccharomyces cerevisiae* strain W303 was used for toxicity assays. *S. cerevisiae* cells were grown at 30°C in YPAD (Yeast extract Peptone Adenine Dextrose) or SD (Synthetic Dextrose minimal) media with agitation, or on YPAD, SD, or SG (Synthetic Galactose minimal) agar plates.

### Plasmid construction and mutagenesis

All cloning procedures were performed by restriction–ligation or restriction-free based methods. Oligonucleotides were purchased from IDT. DNA fragments encoding ARC^tox^ and ARC^imm^ were amplified from *Pantoea ananatis* LMG 20103 using the Q5 polymerase (NEB). Vector backbones and full plasmid templates were amplified using PrimeSTAR Max DNA polymerase (Takara). DNA fragment encoding the *Listeria monocytogenes* LS1292 LXG ARC domain (LisARC) and its cognate immunity was synthesized by Twist Bioscience. PCR products were purified using NucleoSpin Gel and PCR Clean-up columns (Macherey-Nagel), digested with the appropriate restriction enzymes (NEB), and repurified prior to ligation. Ligated plasmids were transformed into *E. coli* DH5α and screened by colony PCR using EconoTaq PLUS GREEN 2x Master Mix (Biosearch Technologies). Plasmid DNA was extracted using either the Wizard Plus SV Minipreps kit (Promega) or the NucleoSpin Plasmid kit (Macherey-Nagel) and verified by Sanger sequencing (Eurofins). Constructs were selected on LB agar plates containing ampicillin (for pRS416_Gal1 and pET-Duet1), chloramphenicol (for pBAD33), kanamycin (for pRSF-Duet1), or streptomycin (for pCDF-Duet1). Glucose (1%) was added for selection with the pNDM220 vector. Toxin and immunity operon were cloned into pET-Duet1, while immunity genes were also cloned individually into pBAD33, pRSF-Duet1, or pCDF-Duet1. Site-directed mutagenesis of ARC^tox^ catalytic residues was performed using QuickChange mutagenesis by PCR-amplifying the whole plasmid using primers bearing the desired mutations and the pNDM220-ARC^tox^ plasmid as template. PCR products were treated by DpnI to eliminate the template plasmid, and phosphorylated and ligated (T4 DNA ligase, NEB) prior to transformation. The pCH10163-ARC^tox^-ARC^imm^ plasmid, encoding ARC^tox^ fused to *E. coli* CdiA was generated by restriction-free cloning, using overlapping regions to fuse the desired fragments.

### Toxicity assays in *E. coli* and yeast

*E. coli* DH5α cells were co-transformed with the pNDM220 and pBAD33 plasmid pairs bearing either ARC^tox^ and ARC^imm^, respectively, or empty vectors as controls. Transformants were selected on LB agar plates supplemented with ampicillin, chloramphenicol, and 1 % of glucose. Overnight cultures grown in the presence of antibiotics and glucose were diluted in 10-fold series, and 10-μL drops were spotted on LB agar plates containing antibiotics and either 1 % glucose (repression conditions) or 1 % arabinose (immunity induction from pBAD33) with 0.05 mM or 0.02 mM IPTG (toxin induction from pNDM220). Plates were incubated at 37 °C for 16-20 hours before imaging.

*S. cerevisiae* strain W303 was transformed with the pRS416_Gal1 encoding ARC^tox^ or an empty vector as control and selected on SD agar plates. Overnight cultures grown in SD liquid medium were diluted in 10-fold series, and 5-μL drops were spotted onto SD (repression condition) and SG (induction condition) agar plates. Plates were incubated at 30 °C for 44-48 hours before imaging.

### Purification of ARC^tox^-ARC^imm^ complex and ARC^tox^

Freshly transformed *E. coli* BL21 (DE3) cells carrying pET-hisTEV-ARC^tox^-ARC^imm^, pRSF-ARC^imm^, or pCDF-ARC^imm^ were grown overnight in LB medium supplemented with appropriate antibiotics. Cultures were diluted 1:100 into 2 L of LB and grown at 37°C to an OD_600_ of 0.8. After addition of 0.5 mM of IPTG, cells were grown overnight at 16 °C with agitation. Cells were collected by centrifugation at 4,000 × g and resuspended in lysis buffer (50 mM Tris-HCl pH 8.5, 250 mM NaCl, 1 mM TCEP) containing cOmplete™ protease inhibitor cocktail (Sigma-Aldrich). Cells were lysed by sonication, and the cell lysate was clarified by centrifugation at 20,000 × g for 45 min. The supernatant was filtered through a 0.45 μm filter and incubated with 2 mL TALON Metal Affinity Resin (Takara) pre-equilibrated with lysis buffer for 1 h at 4 °C with gentle mixing. The resin was washed five times with 15 bed volumes of lysis buffer, and bound proteins were eluted with 1 mL of elution buffer (50 mM Tris-HCl pH 8.5, 250 mM NaCl, 1 mM TCEP, 200 mM imidazole). For purification of ARC toxin alone, after the washing steps with lysis buffer, the resin was incubated with 8 M urea for 2 min, followed by one additional wash with 8 M urea. Refolding was performed by sequential washes with refolding buffer 1 (50 mM Tris-HCl, pH 8.5, 125 mM NaCl, 5% glycerol), followed by two washes with refolding buffer 2 (50 mM Tris-HCl pH 8.5, 250 mM NaCl, 1% glycerol). The toxin was then eluted using 1 mL of elution buffer. The His-TEV tag was removed from both the ARC^tox^/ARC^imm^ complex and ARC^tox^ by incubation with the purified TEV protease (1:100 molar ratio of TEV protease:His-tagged protein) overnight at 4 °C. Proteins were further purified by size-exclusion chromatography using a Superdex 200 Increase 10/300 GL column (Cytiva) equilibrated in buffer A (20 mM Tris-HCl pH 8.5, 150 mM NaCl, 1 mM TCEP) for the ARC^tox^/ARC^imm^ complex, or buffer B (20 mM Tris-HCl pH 8.5, 150 mM NaCl) for ARC^tox^.

### Crystallization, structure determination and refinement of ARC^tox^

Following purification, tag removal, and gel filtration, ARC^tox^ toxin was concentrated to 13.4 mg.mL^-1^ and subjected to crystallization screening using the sitting-drop vapor diffusion method at 20°C. Crystallization drops were set up in Swissci 96-well 2-drop MRC plates (Molecular Dimensions) by mixing 0.5 μL of protein solution with 0.5 μL of reservoir solution. Drops were equilibrated against 70 μL of crystallization screens, including Crystal Screen I and II (Hampton Research), LMB, PACT Premier, JCSG+, and MemGold (Molecular Dimensions). Diffraction-quality crystals of ARC^tox^ were obtained using a reservoir solution containing 0.2 M ammonium chloride, 0.1 M sodium acetate (pH 5.0), and 20% (w/v) PEG 6000. Prior to cryo-cooling in a liquid nitrogen stream, crystals were cryo-protected by briefly transferring them into reservoir solution supplemented with 20% glycerol. X-ray diffraction data were collected at the PROXIMA-2A (PX2A) beamline at the SOLEIL Synchrotron (Gif-sur-Yvette, France), integrated with XDS (Kabsh, 2010) and scaled with AIMLESS (Evans *et al*., 2013) from the CCP4 program suite (Hough *et al.,* 2018). The structure was solved by molecular replacement using a AphaFold3 model of ARC^tox^ as a search probe in PHASER (McCoy *et al*., 2007) with two molecules in the asymmetric unit. The initial map was excellent and the model was improved manually in COOT (Casanal *et al*., 2020) and refined with PHENIX (Adams *et al*., 2010). The final model was refined to a R_work_/R_free_ of 0.18/0.21 with excellent statistics (Supplemental Table S1). The coordinates and structure factors were deposited in the Protein Data Bank (PDB) with accession code 9RO8.

### *In vitro* NAD(P)^+^ hydrolysis assay

Reactions were performed in 100 μL of PBS supplemented with 0.6 mM of NAD^+^ or of NADP^+^ (Roche). Purified, refolded ARC^tox^ was added to a final concentration of 0.1 μM. Samples were incubated at room temperature for 30 min. The reaction was then quenched by adding 50 μL of 6 M NaOH. Fluorescence was measured using TECAN plate reader (excitation: 340 nm; emission: 420 nm) after 30 min of incubation at room temperature in the dark.

### *In vivo* NAD/NADH-Glo assay

*E. coli* strain DH5α cells producing wild-type and mutant ARC^tox^ domains, or both ARC^tox^ and ARC^imm^ were grown in medium supplemented with antibiotics and 1% glucose at 37°C with agitation until OD_600_ reached 0.5. Cells were then washed with LB medium and resuspended in fresh LB supplemented with 0.05 mM of IPTG, 1 % of arabinose, and antibiotics. After incubation for 1 hour at 37°C with agitation, 10^9^ cells were mixed with the NAD/NADH-Glo^TM^ detection reagent (Promega), and the measure of NAD^+^ levels was done according to the manufacturer’s instructions.

### Analysis of ARC^tox^ activity by High Pressure Liquid Chromatography

NAD⁺ or NADP⁺ (1.5 mM) were incubated for 30 min with 0.1 µM of ARC^tox^ or of purified CD38 soluble domain (kind gift of Alain Roussel, LISM, Marseille, France). Proteins were then retained onto Amicon Ultra 3 kDa filters (Sigma-Aldrich) by centrifugation at 11,000 × g for 15 min. The flow-through was injected into an Agilent 1260 Infinity HPLC system equipped with a mixed-mode RP/anion-exchange Atlantis PREMIER BEH C_18_ AX column. Analytes were separated at a flow rate of 1 mL.min^-1^ for 40 min, with a gradient profile of 100% Solvant A (10 mM ammonium formate pH3.0 for NADP^+^, 20 mM ammonium formate pH2.9 for NAD^+^) for 5 min, linear gradient to 50% Solvant A / 50% Solvent B (acetonitrile) for 30 min, and 100% Solvent B for 5 min. Analytes were detected by absorbance at 254 nm. The retention times were compared to standards run on the same column (Nicotinamide (NAM) 250-450 µM, NAD⁺ 40 µM, NADP⁺ 150 µM, ADP-ribose 150-200 µM, cyclic ADP-ribose 60-160 µM)

### Bacterial competition assay

Interbacterial competition was measured using the LAGA assay (Taillefer et al., 2023). This assay is based on the degradation of chlorophenol-red β-D-galactopyranoside (CPRG), a membrane-impermeable chromogenic substrate of the β -galactosidase, which is cleaved by the β -galactosidase released by the recipient cell lysis. *E. coli* DH5α cells producing CdiB and the CdiA chimera proteins were used as donors whereas *E. coli* W3110 (*lacZ*^+^) cells were used as recipient. Both strains were grown separately to an OD_600_ of 0.5, washed twice with LB medium supplemented with 0.05 mM IPTG, and mixed to a 1:2 (donor:recipient) ratio. Ten µL of the mixtures were spotted onto LB agar plates supplemented with 0.05 mM of IPTG. Plates were incubated for 4 hours at 37 °C, and recipient cell lysis was observed by the addition of a 10-µL drop of 2 mM of CPRG (Roche) on the top of each spot.

### Statistical analysis

*In vitro* ARC^tox^ activity data were analyzed using an unpaired *t*-test. *In vivo* data from NAD/NADH-Glo^TM^ assay were analyzed using one-way ANOVA followed by Dunnett’s post hoc test. All analyses were performed using the GraphPad Prism software.

## DATA AVAILABILITY

The *Pantoea ananatis* PANA_2924 ARC^tox^ structure has been deposited to the Protein Data Bank (PDB) under accession code 9RO8.

## SUPPORTING INFORMATION

This article contains supporting information (two Supplemental Figures and four Supplemental Tables).

## ACKNOWLEDGMENTS

We thank the members of the Cascales and Jurėnas laboratories for discussions and support, Vincent Géli, Pierre Luciano and Frédéric Jourquin (CRCM, Marseille, France) for providing yeast strains, vectors and protocols, Alain Roussel (LISM, Marseille, France) for the kind gift of purified human CD38, the staff at the SOLEIL PROXIMA 2 beamline, and Audrey Gozzi for technical support.

## LEGENDS TO SUPPLEMENTAL FIGURES

**Supplemental Figure S1.**
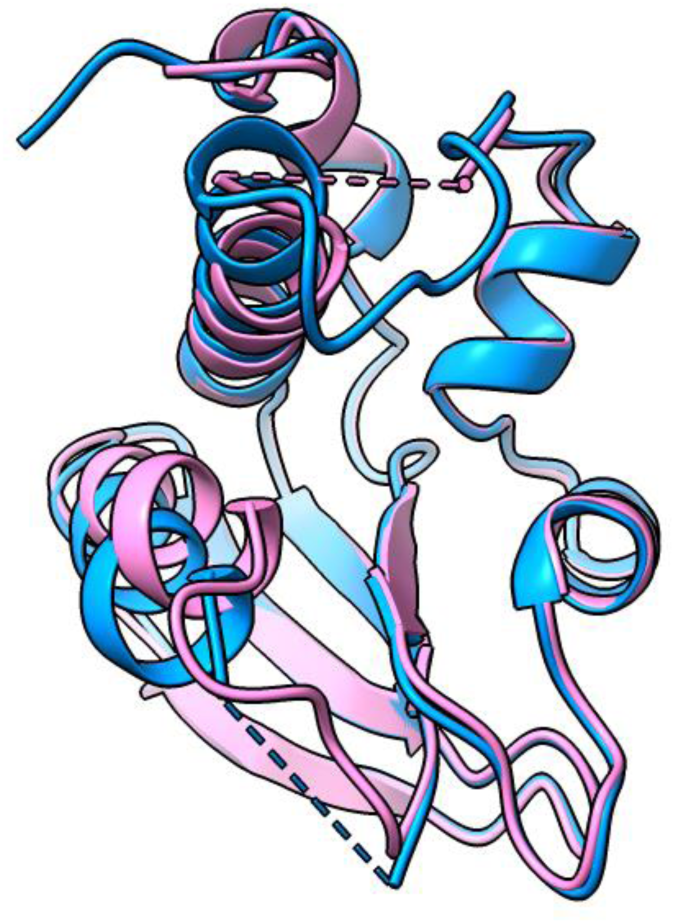
Comparison of the crystal structures of the two chains of the PANA_2924 C-terminal extension asymmetric unit. Superimposition of the crystal structures of the chains A (pink) and B (blue) of the PANA_2924 C-terminal domain. The dotted lines highlight the positions of the loops that are missing in one structure but visible in the other.

**Supplemental Figure S2.**
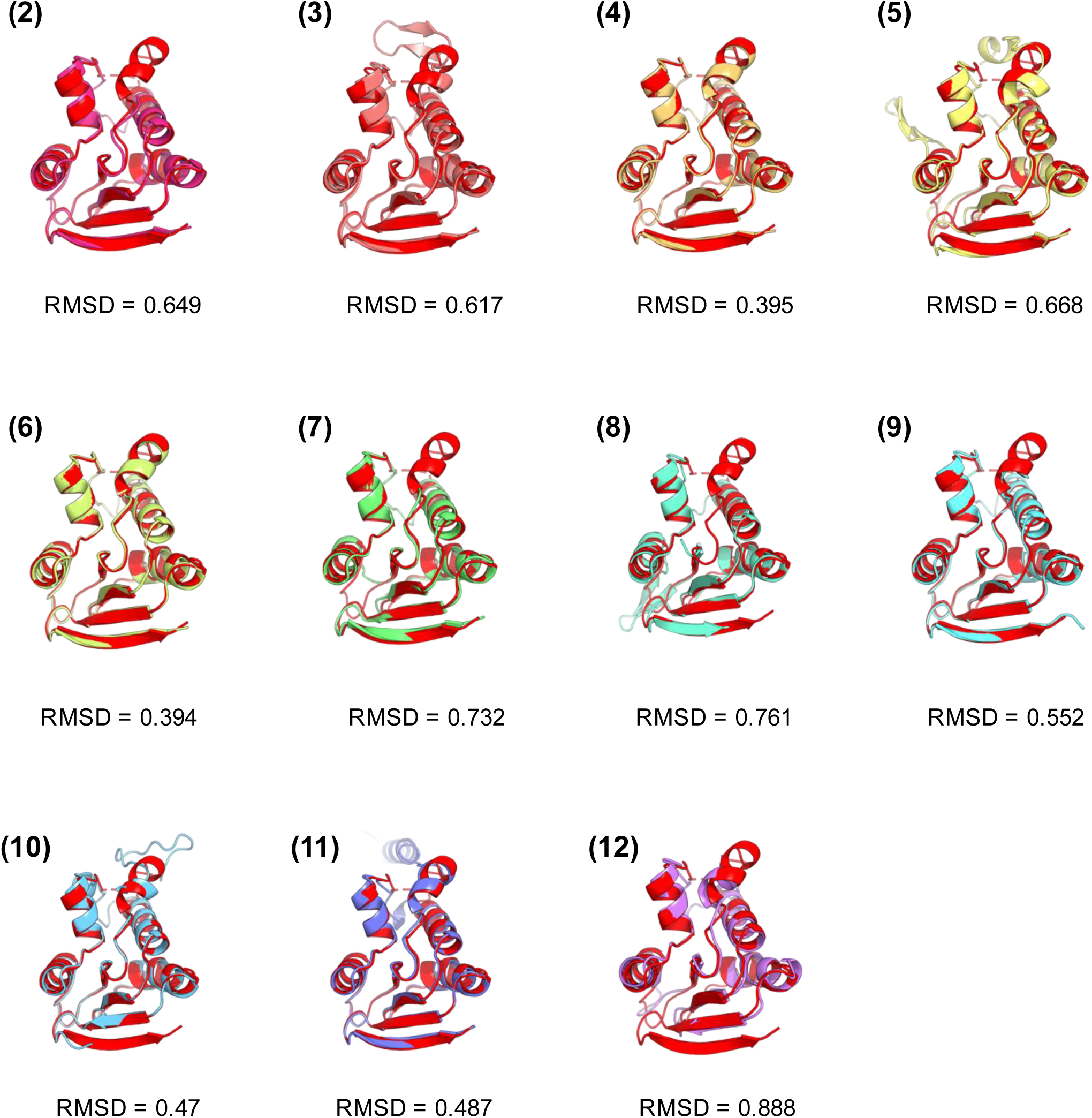
AlphaFold3 structural models of ARC domains (2, *Agrobacterium rubi* NTF28035; 3, *Burkholderia gladioli* WP_186165364; 4, *Listeria monocytogenes* WP_120135998; 5, *Gordonia jinhuaensis* WP_188589035; *Lactiplantibacillus plantarum* WP_076633504; 7, *Stenotrophomonas maltophilia* OCK46403; 8, *Nonomuraea fuscirosea* WP_364663299; 9, *Marinobacter shengliensis* WP_138437386; 10, *Pendulispora albinea* WP_394825747; 11, *Allomuricauda sp.* RPG31737; 12, *Streptomyces mirabilis* WP_388533272) aligned with the *P. ananatis* ARC^tox^ crystal structure (red). Root-Mean-Square Deviation (RMSD) values are indicated below each superimposition.

---

**Supplemental Table 1.** Data collection and refinement statistics.

|  | **ARC<sup>tox</sup> (PDB : 9RO8)** |
| --- | --- |
| **Data collection** |  |
| **Resolution range** | 39.66 - 1.594 (1.63 - 1.59) |
| **Space group** | P 21 21 21 |
| **Unit cell** | 44.23 70.213 96.119 90 90 90 |
| **Total reflections** |  |
| **Unique reflections** | 40577 (2746) |
| **Multiplicity** |  |
| **Completeness (%)** | 99.73 (96.38) |
| **Mean I/sigma(I)** | 17.0 (2.2) |
| **Wilson B-factor** | 21.48 |
| **R-merge** | 0.092 (1.053) |
| **R-meas** | 0.096 (1.099) |
| **R-pim** | 0.027 (0.31) |
| **CC1/2** | 0.99 (0.76) |
| **Refinement** |  |
| Reflections used in refinement | 40577 (2746) |
| Reflections used for R-free | 2000 (136) |
| R-work | 0.1826 (0.2430) |
| R-free | 0.2145 (0.2877) |
| Number of non-hydrogen atoms | 2308 |
| macromolecules | 2016 |
| ligands | 0 |
| solvent | 292 |
| Protein residues | 258 |
| RMS(bonds) | 0.010 |
| RMS(angles) | 1.04 |
| Ramachandran favored (%) | 100.00 |
| Ramachandran allowed (%) | 0.00 |
| Ramachandran outliers (%) | 0.00 |
| Rotamer outliers (%) | 0.45 |
| Clashscore | 1.74 |
| Average B-factor | 27.34 |
| macromolecules | 26.40 |
| solvent | 33.83 |

Statistics for the highest-resolution shell are shown in parentheses.

**Supplemental Table S2.**
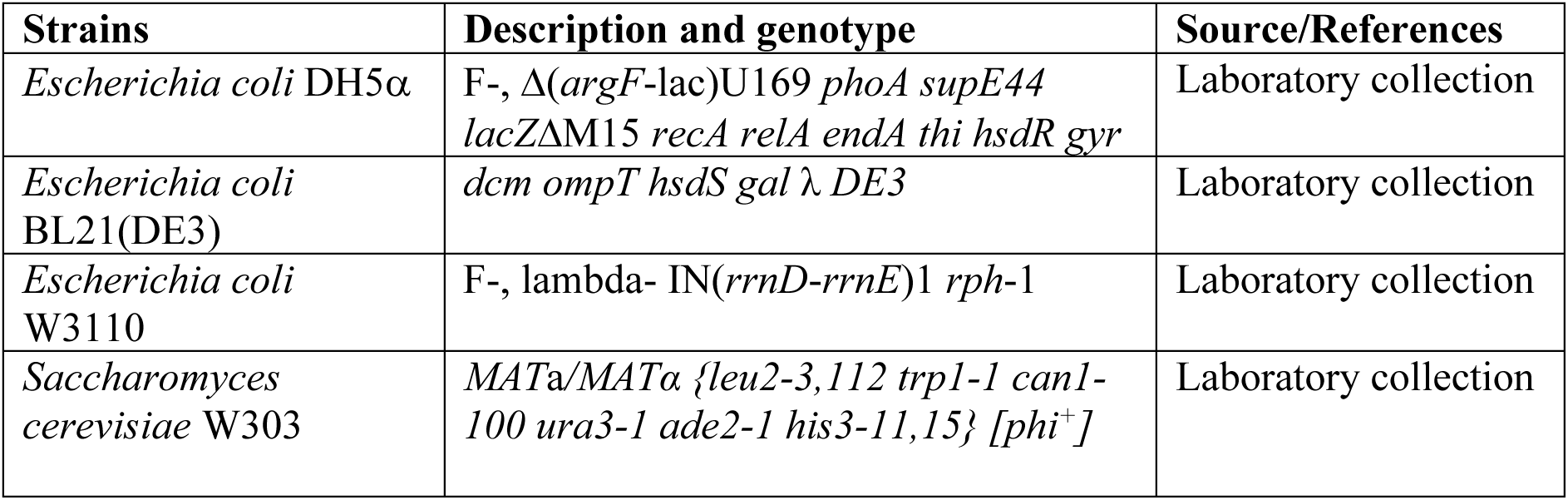
Strains used in this study.

**Supplemental Table S3.**
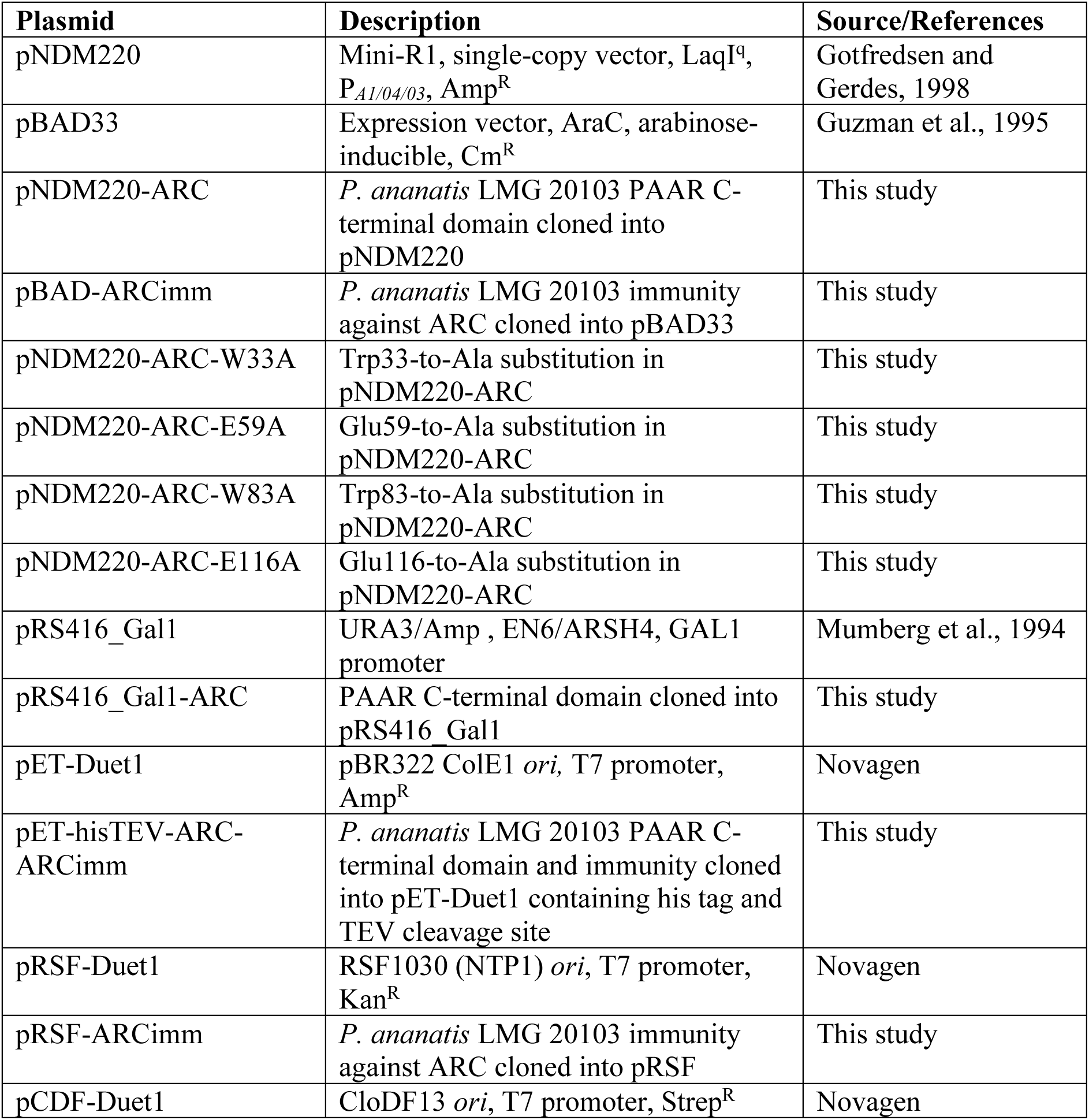
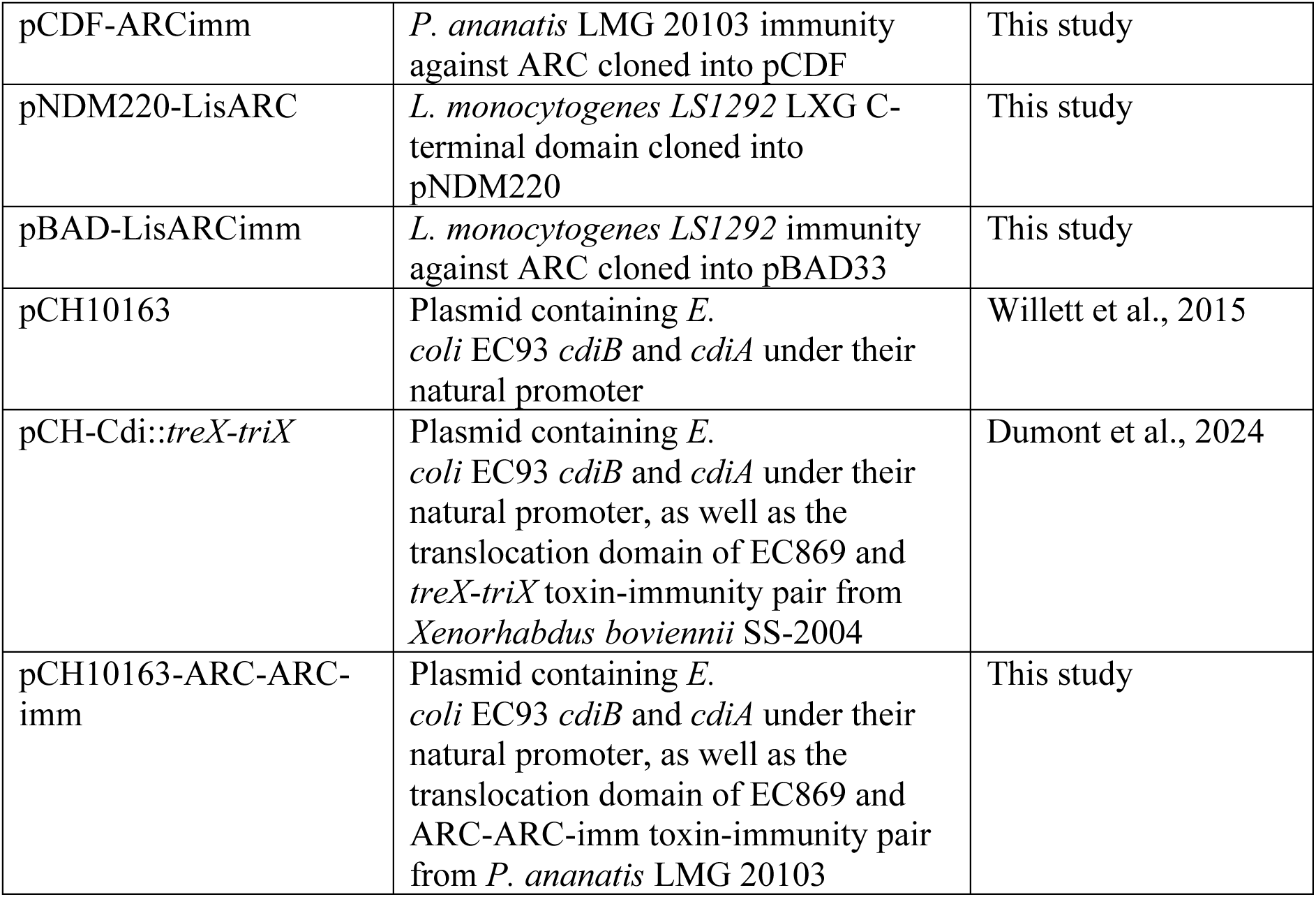
Plasmids used in this study.

**Supplemental Table S4.**
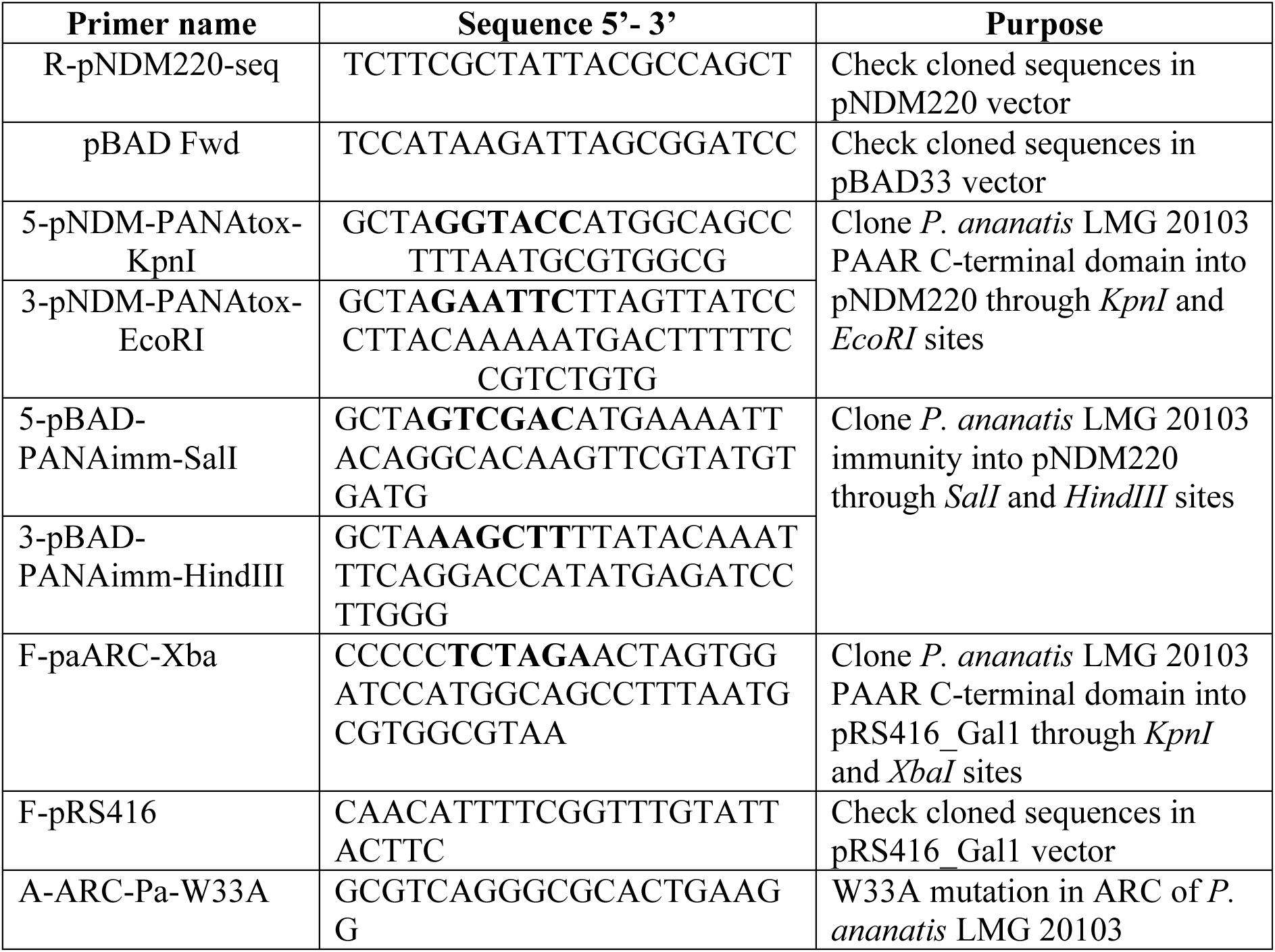
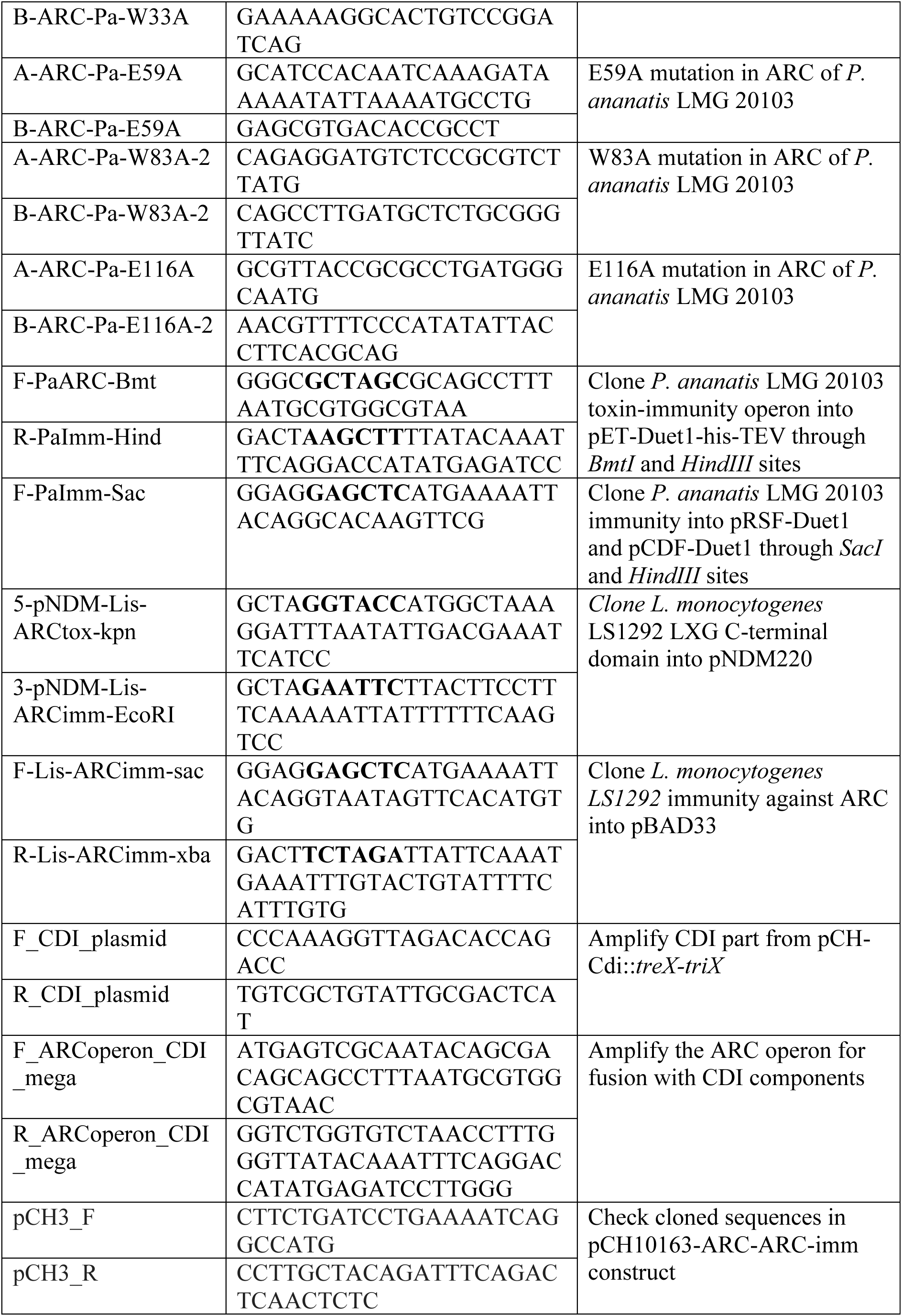
Primers used in this study.

